# Investigation of the global translational response to oxidative stress in the model archaeon *Haloferax volcanii* reveals untranslated small RNAs with ribosome occupancy

**DOI:** 10.1101/2025.04.08.647799

**Authors:** Emma Dallon, Haley M. Moran, Sadhana R. Chidambaran, Arman Kian, Betty Y.H. Huang, Stephen D. Fried, Jocelyne DiRuggiero

**Affiliations:** Department of Biology, Johns Hopkins University, Baltimore, Maryland, USA; Department of Chemistry, Johns Hopkins University, Baltimore, Maryland, USA; T. C. Jenkins Department of Biophysics, Johns Hopkins University, Baltimore, Maryland, USA; Department of Earth and Planetary Sciences, Johns Hopkins University, Baltimore, Maryland, USA

## Abstract

Oxidative stress induces a wide range of cellular damage, often causing disease and cell death. While many organisms are susceptible to the effects of oxidative stress, haloarchaea have adapted to be highly resistant. Several aspects of the haloarchaeal oxidative stress response have been characterized, however little is known about the impacts of oxidative stress at the translation level. Using the model archaeon *Haloferax volcanii*, we performed RNA-seq and ribosome profiling (Ribo-seq) to characterize the global translation landscape during oxidative stress. We identified 281 genes with differential translation efficiency (TE). Downregulated genes were enriched in ribosomal and translation proteins, in addition to peroxidases and genes involved in the TCA cycle. We also identified 42 small noncoding RNAs (sRNAs) with ribosome occupancy. Size distributions of ribosome footprints revealed distinct patterns for coding and noncoding genes, with 12 sRNAs matching the pattern of coding genes, and mass spectrometry confirming the presence of seven small proteins encoded in these sRNAs. However, the majority of sRNAs with ribosome occupancy had no evidence of coding potential. Of these ribosome-associated sRNAs, 12 had differential ribosome occupancy or TE during oxidative stress, suggesting that they may play a regulatory role during the oxidative stress response. Our findings on ribosomal regulation during oxidative stress, coupled with potential roles for ribosome-associated noncoding sRNAs and sRNA-derived small proteins in *H. volcanii*, revealed additional regulatory layers and underscore the multifaceted architecture of stress-responsive regulatory networks.

**Importance:** Archaea are found in diverse environments, including as members of the human microbiome, and are known to play essential ecological roles in major geochemical cycles. The study of archaeal biology has expanded our understanding of the evolution of eukaryotes, uncovered novel biological systems, and revealed new opportunities for applications in biotechnology and bioremediation. Many archaeal systems, however, remain poorly characterized. Using *Haloferax volcanii* as a model, we investigated the global translation landscape during oxidative stress. Our findings expand current knowledge of translational regulation in archaea and further illustrate the complexity of stress-responsive gene regulation.

## Introduction

Since the advent of oxygenic metabolism, oxidative stress has affected most organisms across all domains of life. This stress is associated with reactive oxygen species (ROS) that disrupt cellular redox state and covalently damage essential macromolecules (1–3). While endogenous ROS, including hydrogen peroxide and superoxide, arise as byproducts of oxygen metabolism (1), ROS levels can increase due to environmental stressors (1, 4). Prolonged oxidative stress induces a range of responses, including disease, autophagy, and cell death (5–8). To neutralize ROS, prokaryotes utilize numerous enzymes (9–13), and certain bacteria and archaea, such as haloarchaea, are known to be exceptionally resistant to oxidative stress (4, 14–17).

Archaea have attracted interest because of mounting evidence that they are ancestral to eukaryotes (18). This is reflected in the mosaic nature of archaeal cells with prokaryotic features and yet information processing machinery more closely related to that of eukaryotes (18–20). The study of archaea has led to the discovery of novel biological systems, such as unique lipids and protein modifications not found in other domains (21, 22). These findings shed light on the evolution of eukaryotes from a prokaryotic ancestor (23) and expand possibilities for biotechnology applications (24). However, many archaeal regulatory systems remain poorly characterized, and our knowledge of them lags behind that of bacterial and eukaryotic counterparts. For example, mechanisms of transcriptional regulation, especially in response to oxidative stress, have only been studied in a few archaeal species (25–27). In the model archaeon, *Haloferax volcanii*, a key transcriptional regulator, RosR, was shown to affect the expression of many genes in response to oxidative stress (26). However, while RosR is conserved among haloarchaea, its function varies among species (25). Gene regulation at the post-transcriptional level by small noncoding RNAs (sRNAs) is particularly critical to stress responses in many prokaryotic organisms (28–31), and has been linked to oxidative stress in several bacteria (32–34) and in the archaeon *H. volcanii* (27, 35). In *H. volcanii*, the most upregulated sRNA during oxidative stress (SHOxi) binds and destabilizes malic enzyme transcripts, leading to higher cell survival rates during oxidative stress (35).

While many studies have characterized mechanisms of oxidative stress response at the transcriptional, post-transcriptional, and post-translational levels, little is known about the impacts of oxidative stress at the translation level. Translational regulation has been shown to play a major role in various stress conditions in both bacteria and yeast (36–39). In yeast, the global translation landscape during oxidative stress has been explored using ribosome profiling (Ribo-seq), revealing the regulation of several individual genes and general shifts in translation patterns across all genes (40). Mechanisms of translational regulation by proteins and sRNAs during stress have also been reported in archaea. The archaeal protein IF6, a homolog to eukaryotic IF6, inhibits translation by binding to the large ribosomal subunit, preventing assembly of the full ribosome, and is overexpressed during thermal stress (41). In *Methanosarcina mazei*, sRNAs were shown to mask the ribosome binding site of target mRNAs to modulate translation in response to nitrogen starvation (42, 43). This mechanism bears resemblance to an oxidative stress-responsive sRNA in *E. coli* (44). Additionally, proteomic studies in *H. volcanii* have begun to elucidate the differences between transcription and translation during stress responses (45, 46). However a comprehensive study of translational regulation during oxidative stress response has thus far been lacking.

In this work we present evidence of translational regulation during oxidative stress in *H. volcanii*. This model archaeon is well-characterized at the genetic level (47–49) and a variety of genetic and biochemical tools are available for its manipulation (50, 51). Recent advances in translation research in *H. volcanii* (52, 53) have demonstrated that Ribo-seq enables the detection of translational changes, providing a means to probe the impact of oxidative stress on protein synthesis. In the present work, we combined RNA sequencing (RNA-seq) and ribosome profiling (Ribo-seq) to determine the global translation landscape during oxidative stress response. We identified genes that are differentially regulated at the translation level under oxidative stress and, using an integrative approach, investigated the potential role of stress-responsive ribosome-associated sRNAs and sRNA-encoded small proteins.

## Results

We investigated the global translation landscape in *H. volcanii* during oxidative stress by performing RNA-seq and Ribo-seq *on H. volcanii* cultures with and without stress treatment. Ribo-seq, a technique that captures ribosome occupancy across thousands of transcripts, can be used as a proxy for translation (54). By comparing transcription levels from RNA-seq data to ribosome occupancy from Ribo-seq data, we assessed differential expression, ribosome occupancy, and translation efficiency, allowing us to identify genes regulated at the translation level. Using this integrated approach, we also identified ribosome-associated sRNAs responsive to oxidative stress.

### Hundreds of genes, including known oxidative stress-responsive genes, had differential ribosome occupancy during oxidative stress

Libraries for RNA-seq and Ribo-seq were prepared from four biological replicate cultures treated with 2 mM hydrogen peroxide for one hour to induce oxidative stress, as previously described (27, 35). Four replicates without stress treatment were used as controls. Sequenced reads were aligned to the genome (49) and normalized expression values of transcripts per million (TPM) were calculated (Table S1, Table S2). Using a minimum detection cutoff of 10 TPM per gene, averaged across all biological replicates, we detected 84% of annotated genes in both RNA-seq control and stress samples. For the Ribo-seq data, we detected 74% of annotated genes in control samples and 80% in stress samples.

We calculated differential expression between controls and oxidative stress treatments for the RNA-seq and Ribo-seq datasets (Table S3). Genes with an adjusted p-value <0.05 and an absolute log_2_ fold change >1 were considered differentially expressed during oxidative stress. Lowly expressed genes with TPM <10 in both conditions were excluded from the analysis. We found 559 genes with differential RNA expression during oxidative stress, in contrast to 966 genes with differential ribosome occupancy (Fig. 1A, B).

**Figure 1.**
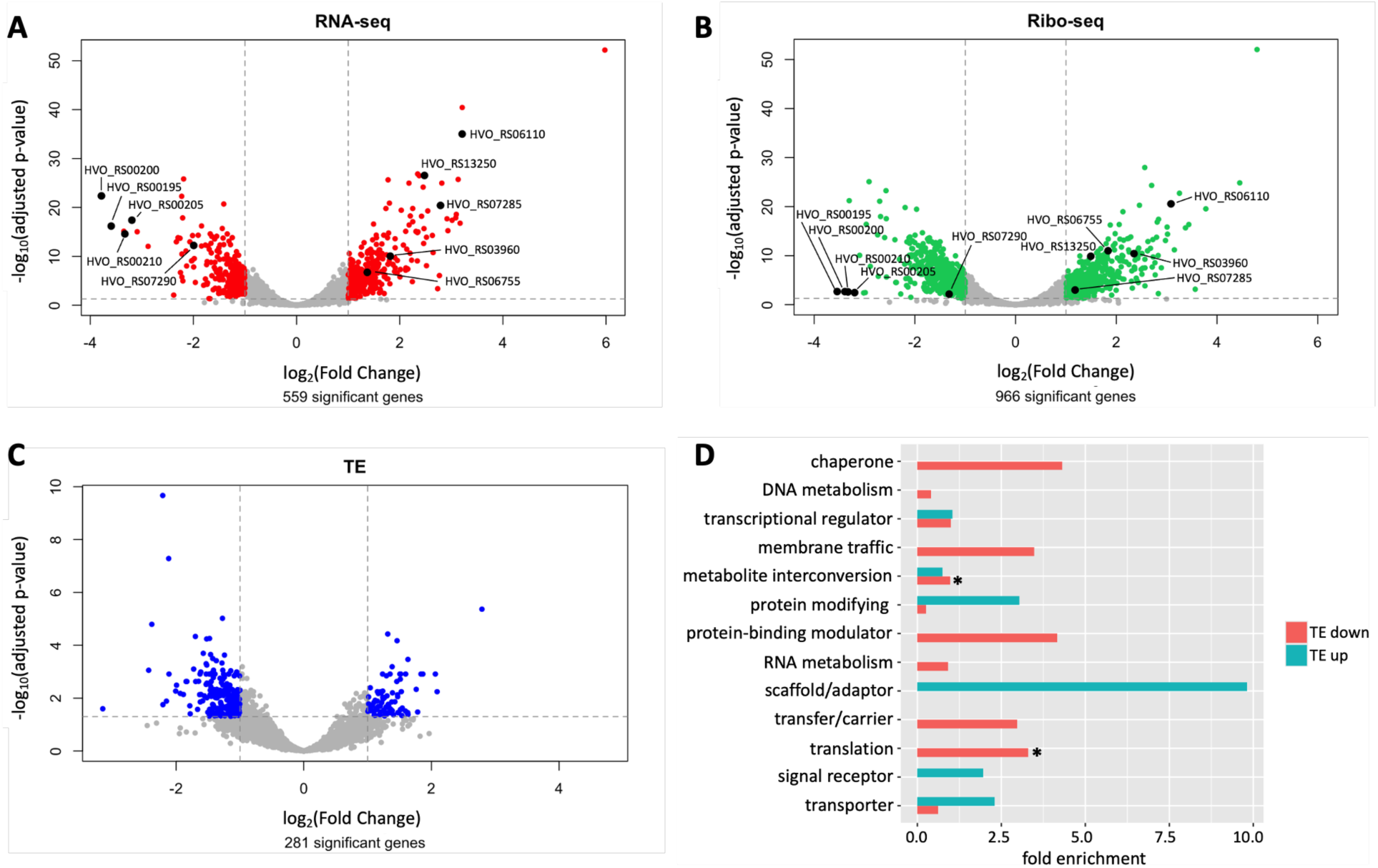
Comparison of genes with differential transcription, ribosome occupancy, and translation during oxidative stress. Volcano plots of log_2_ fold changes and corresponding -log_10_ p-values of transcript abundances (A), ribosome occupancies (B), and translation efficiency (C) comparing stress vs no stress conditions. Red, green, and blue colored points highlight transcripts with differential expression (adjusted p-value <0.05 and absolute log_2_ fold change >1). Fold change and p-values were calculated using DESeq2. Labeled black points indicate genes previously shown to be involved in oxidative stress response. All other genes are shown in gray. D. Functional categories (protein class) of genes with differential TE during oxidative stress. Functions of significant genes were categorized using PANTHER generic mapping based on homology. Enrichment of functional categories in upregulated (TE up) or downregulated (TE down) genes compared to all annotated genes was calculated using the total number of PANTHER hits for each group (2061 for all annotated genes, 99 for TE down, and 42 for TE up). Asterisks indicate significant categories (translation) or categories that contain significantly enriched subcategories (ribosomal proteins; peroxidases).

To validate our experimental approach, we investigated the differential expression of five previously known oxidative stress-responsive genes. These genes were for superoxide dismutase (HVO_RS03960), catalase (HVO_RS13250), a universal stress protein (HVO_RS06755), and a ssDNA binding replication protein (HVO_RS06110); the latter two were both reported as upregulated proteins in *H. volcanii* under hypochlorite stress (46). All showed significant upregulation in both RNA-seq and Ribo-seq datasets (Fig1A, B; Table S4), implying our experimental approach recapitulated known aspects of oxidative stress response. We also observed upregulation of a gene involved in metal homeostasis (HVO_RS07285; Fig1A, B; Table S4) and downregulation of four others (HVO_RS00195-210; HVO_RS07290; Fig1A, B; Table S4), consistent with previous findings (46).

We then tested the hypothesis that genes with differential transcript expression would have differential ribosome occupancy, as transcripts highly abundant during oxidative stress would likely be bound by a higher number of ribosomes than transcripts with low abundance. By comparing significant gene lists from RNA-seq and Ribo-seq data, we identified 320 genes with both differential RNA expression and ribosome occupancy during stress. The remaining 646 genes changed only in ribosome occupancy during stress (67% of all genes with differential ribosome occupancy; Fig. S1). Several genes had differential ribosome occupancy in accordance with previously published proteomics data [HVO_RS02870-*msrA;* HVO_RS04695-cysteine desulfurase; Table S3; (46)]. Overall, in both RNA-seq and Ribo-seq datasets, we detected the majority of annotated genes and identified hundreds of genes with differential ribosome occupancy during oxidative stress.

### Translation efficiency revealed global downregulation of translation during oxidative stress

To increase the stringency of our analysis and identify genes that were regulated uniquely at the translation level, we calculated translation efficiency (TE). This quantity is defined as the ratio between the Ribo-seq fold change and the RNA-seq fold change and was used for differential expression analysis (Table S3). TE values account for changes in transcript availability and identify genes that change at the Ribo-seq and/or RNA-seq level while eliminating those that change proportionally in both. We found 281 genes with significant differential TE (Fig. 1C; adjusted p-value <0.05, absolute log_2_ fold change >1). When comparing genes with differential TE to those with differential transcript expression and differential ribosome occupancy, we found that 240 genes (85.4%) were regulated only at the level of translation (Fig. S2). The remaining 41 genes were regulated at both the transcriptional and translational levels, with 30 genes displaying opposing effects at each level (Table S3). The majority (66.7%) of genes with opposing effects were upregulated at the transcriptional level yet downregulated at the translational level. Of the 281 genes with differential TE, 29.9% were upregulated and 70.1% were downregulated during oxidative stress (Table S5).

We investigated some basic properties of transcripts with differential TE to determine whether they were distinct from the overall transcriptome. We found no differences in amino acid or codon frequencies (Fig. S3) and no difference in the proportion of leaderless transcripts compared to the whole transcriptome (Table S5), most likely because *H. volcanii* translation initiation is not dependent on the presence of a 5’UTR (55). The average size for genes with differential TE was slightly smaller, likely due to a narrower size range of significant genes (Table S5).

We next investigated whether genes with differential TE during oxidative stress exhibited functional enrichments. Gene ontology (GO) analysis was performed using PANTHER generic mapping (56, 57), with enrichment significance determined via Fisher’s test and false discovery rate (FDR correction). Additionally, KEGG pathway enrichment analysis was conducted using the STRING database (58). Both PANTHER and STRING rely on homology-based functional predictions. Approximately half of all annotated *H. volcanii* genes were assigned to functional categories using PANTHER (Table S6). However, few significant functional enrichments were observed among genes with differential TE, likely due to small sample size (Fig. 1D, Table S7). Among genes with downregulated TE during oxidative stress, we observed significant enrichment of translation-related functions (Fig. 1D). Specifically, GO enrichment analysis revealed overrepresentation of ribosomal proteins (fold enrichment = 4.41, FDR = 0.0005) and translation-associated proteins (fold enrichment = 3.39, FDR = 0.0008). Consistent with these results, KEGG pathway analysis also identified significant enrichment of ribosomal proteins (FDR = 0.00093). In addition, we observed differential TE for many genes associated with redox homeostasis. Notably, downregulated genes showed enrichment of peroxidases (GO; fold enrichment = 9.5, FDR = 0.0213) and tricarboxylic acid (TCA) cycle components (KEGG; FDR = 0.0242). In contrast, no statistically significant functional enrichment was detected among genes with upregulated TE during oxidative stress (Fig. 1D).

In addition to analyzing functional categories of genes with differential TE, we sought to identify whether specific cellular pathways were subject to translational regulation during oxidative stress. To do so, we used the STRING database to predict interaction networks for genes with upregulated or downregulated TE. Few predicted interactions were identified among upregulated genes (Fig. S4A). In contrast, genes with downregulated TE formed a larger, interconnected network with many functional clusters (Fig. S4B). The two largest clusters were composed of ribosome-associated genes, including numerous ribosomal proteins, and genes involved in the TCA cycle. Additional smaller clusters, with four genes each, were related to oxidative phosphorylation, pyrimidine biosynthesis, and putative translation initiation factors. These interaction networks suggest that while global translation is broadly reduced under oxidative stress, specific pathways are subject to targeted translational regulation, likely to modulate metabolic activity and maintain cellular homeostasis.

While the majority of genes with differential TE were regulated only at the translational level, 15% of genes with differential TE exhibited changes (absolute log_2_ fold change >1) at both the transcriptional and translational levels. One such gene, HVO_RS04895, encoding a protein with a transmembrane domain, showed increased ribosome occupancy and TE (Table S8). This gene is known to be transcriptionally upregulated during oxidative stress via the transcriptional regulator OxsR (59). We also identified two genes with a distinct regulation pattern characterized by downregulated TE, coupled with increased RNA expression and ribosome occupancy (Table S8). These genes encode KatG (HVO_RS13250), a catalase/peroxidase that uses hydrogen peroxide as its substrate (60, 61), and DpsA (HVO_RS07285), a DNA starvation/stationary phase protection protein (62, 63). Previous proteomics studies reported no significant change in KatG protein levels, while DpsA was upregulated during oxidative stress (46). These findings suggest that transcriptional changes alone may not fully predict protein abundance under stress conditions, and that translational regulation likely serves as an additional rapid mechanism for fine-tuning protein levels during stress.

### Many small noncoding RNAs had ribosome occupancy

As expected, most ribosome footprints in our dataset corresponded to annotated coding sequences. However, it was remarkable that several small noncoding RNAs (sRNAs), previously identified by sRNA-seq (27), also exhibited ribosome occupancy. In total, 42 out of 56 sRNAs detected in our RNA-seq dataset showed ribosome occupancy (TPM >10; Table S9), with 17 of these sRNAs displaying differential ribosome occupancy or TE during oxidative stress (Table 3).

To investigate the coding potential of sRNAs with ribosome occupancy, we used several computational approaches to identify putative small ORFs (smORFs; Fig. 2A). Public database searches revealed nine annotated smORFs encoded by sRNAs (Fig. 2A). Of these, three have been detected by mass spectrometry (MS) as reported in the Archaeal Proteome Project database (64). Next, we used REPARATION (65), a tool designed to predict novel ORFs from ribosome profiling data (53). Using our ribosome profiling dataset, REPARATION accurately predicted 82% of all annotated proteins (Table S10), including 436 of 542 (80%) annotated smORFs (≤100 amino acids). In addition, REPARATION predicted 194 novel genes (Table S11) of which only seven encoded small proteins and one originated from a ribosome-associated sRNA (Fig. 2A). While REPARATION predicted few smORFs within sRNAs, sequence-based tools such as ORFfinder were overly permissive [Fig. 2A; (66)], often predicting ORFs in sRNAs even in the absence of a stop codon within or around the sRNA sequence (Table S12).

**Figure 2.**
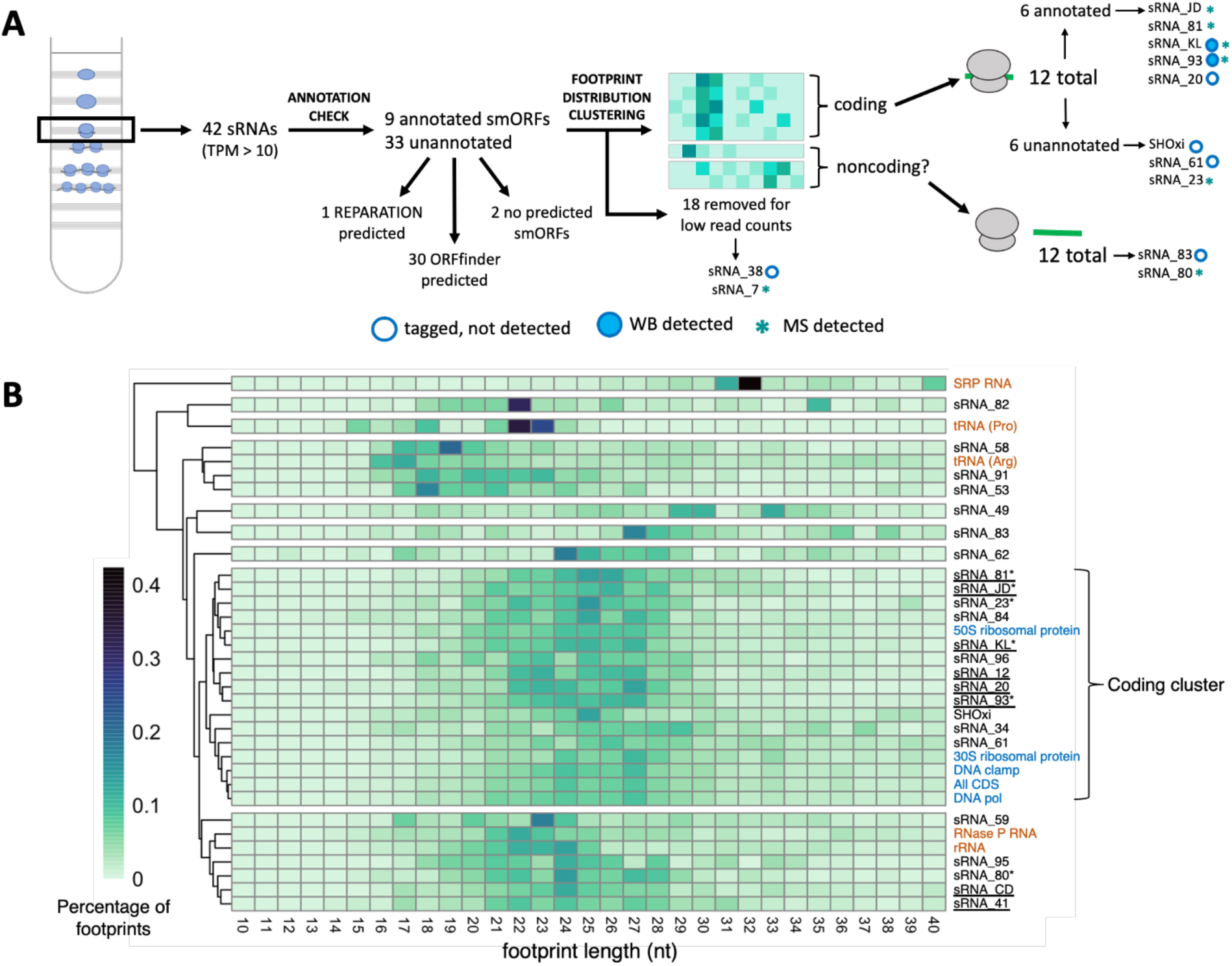
Ribosome footprint length distributions and sRNAs with ribosome occupancy. A. Schematic of analyses to assess the coding potential for sRNAs with ribosome occupancy, starting with the isolation of monosome fractions during ribosome profiling. B. Clustered heatmap of ribosome footprint length distributions for sRNAs, coding genes, and noncoding genes with ribosome occupancy under control conditions. Darker colors indicate higher read percentage. Known coding genes are shown in blue, known noncoding genes in orange, and sRNAs in black. sRNAs with annotated proteins are underlined. sRNAs for which peptides have been detected by MS are marked with an asterisk. Percentages for ‘All CDS’ were calculated using sum read counts of all coding genes.

These conflicting computational predictions, combined with empirical evidence for coding potential for only 10 out of 42 sRNAs with ribosome occupancy (Fig. 2A), prompted further investigation of the nature of the ribosome occupancy on sRNAs. To distinguish between true translation and non-canonical ribosome association, we examined the ribosome footprint length distribution. Footprint length distributions have previously been used to infer translation states at a genome-wide level (52), and we applied this approach to individual sRNAs to assess their coding potential.

### Size distributions of ribosome footprints revealed distinct patterns for coding and noncoding genes

We analyzed ribosome footprint length distributions of coding genes, noncoding genes, and sRNAs to assess the coding potential of ribosome-associated sRNAs. As expected, the distribution of footprint lengths for coding genes - both at the level of individual genes - and across the sum of all coding genes - was sharply enriched in footprints between 24 and 27 nucleotides (nt) in length (Fig. 2B). This was consistent with previous work identifying that the characteristic footprint length for elongating ribosomes was 27 nt (52). In contrast, noncoding genes, including rRNAs, SRP RNA, RNaseP RNA, and tRNAs, displayed distinct footprint length distributions that were markedly different from those of coding genes (Fig. 2B).

To further explore sRNA coding potential, we performed clustering analysis based on the footprint length distributions of coding genes, noncoding genes, and sRNAs of interest. Clustering was optimized using the minimum number of clusters required to separate known coding genes from rRNAs and tRNAs. When clustering sRNAs with annotated smORFs (Fig. S5A), we observed that sRNAs with low total read counts did not consistently group with coding genes (Fig. S5B), indicating that sufficient read count is necessary to accurately infer coding potential. From this analysis, we determined that a minimum of 65 read counts across all biological replicates was required for reliable classification.

Using this threshold value, we found that 12 sRNAs clustered with known coding genes based on their footprint length distribution (Fig. 2B), seven of which corresponded to sRNAs with annotated smORFs. The high proportion of sRNAs with annotated smORFs included in the coding cluster supports the utility of footprint length distribution as a means of distinguishing coding from noncoding sRNAs, providing sufficient read coverage is available. However, we also identified 12 sRNAs ribosome-associated sRNAs that lacked both annotated smORFs and footprint length distributions consistent with coding genes.

### Discovery of small proteins encoded by sRNA transcripts

To validate predictions of coding sRNAs and assess whether predicted small proteins encoded by sRNAs could be detected, we used liquid chromatography tandem mass spectrometry (LC-MS/MS). This analysis included three biological replicates with and without oxidative stress treatment, the latter chosen to induce the expression of stress-responsive sRNAs. We sequentially passed cell lysates through centrifugal filters with 100 kDa and 30 kDa molecular cutoffs to deplete high molecular weight proteins (Fig. S6).

Our label-free quantitative differential expression analysis detected 2,095 proteins, filtered to 5% false discovery rate (FDR) and with at least two peptide spectrum matches (PSMs). This dataset represented 53% of the total proteome of *H. volcanii*, with a median protein sequence coverage of 59%, consistent with other high-quality MS datasets reported for this organism (64). As previously reported (67), centrifugal filtering is an imperfect method to enrich for small proteins, and indeed, the size distribution of the proteins we detected closely represented that of the full proteome (Fig. 3A). Notably, we did not detect any protein smaller than 25 amino acids, despite the existence of predicted smORFs as short as 10 amino acids (Fig. 3B). Nevertheless, we detected 235 smORFs (Fig. 3B) ranging from 31 to 100 amino acids in length, representing 40% of annotated smORFs (Table S13). Among these, seven smORFs were encoded within putative sRNAs, including four previously annotated (Fig. 2A; Table 1). For five out of these seven sRNA-encoded proteins, ribosome footprint length distributions clustered with those of known coding genes (Fig. 2B) and two smORFs, sRNA_93 and sRNA_KL, were highly conserved among haloarchaea. However, computational predictions, including sequence and structural homology searches, did not reveal functional annotations for these smORFs. Despite these findings, seven sRNAs with footprint length distributions consistent with coding genes lacked detectable protein-level evidence in our MS dataset.

**Figure 3.**
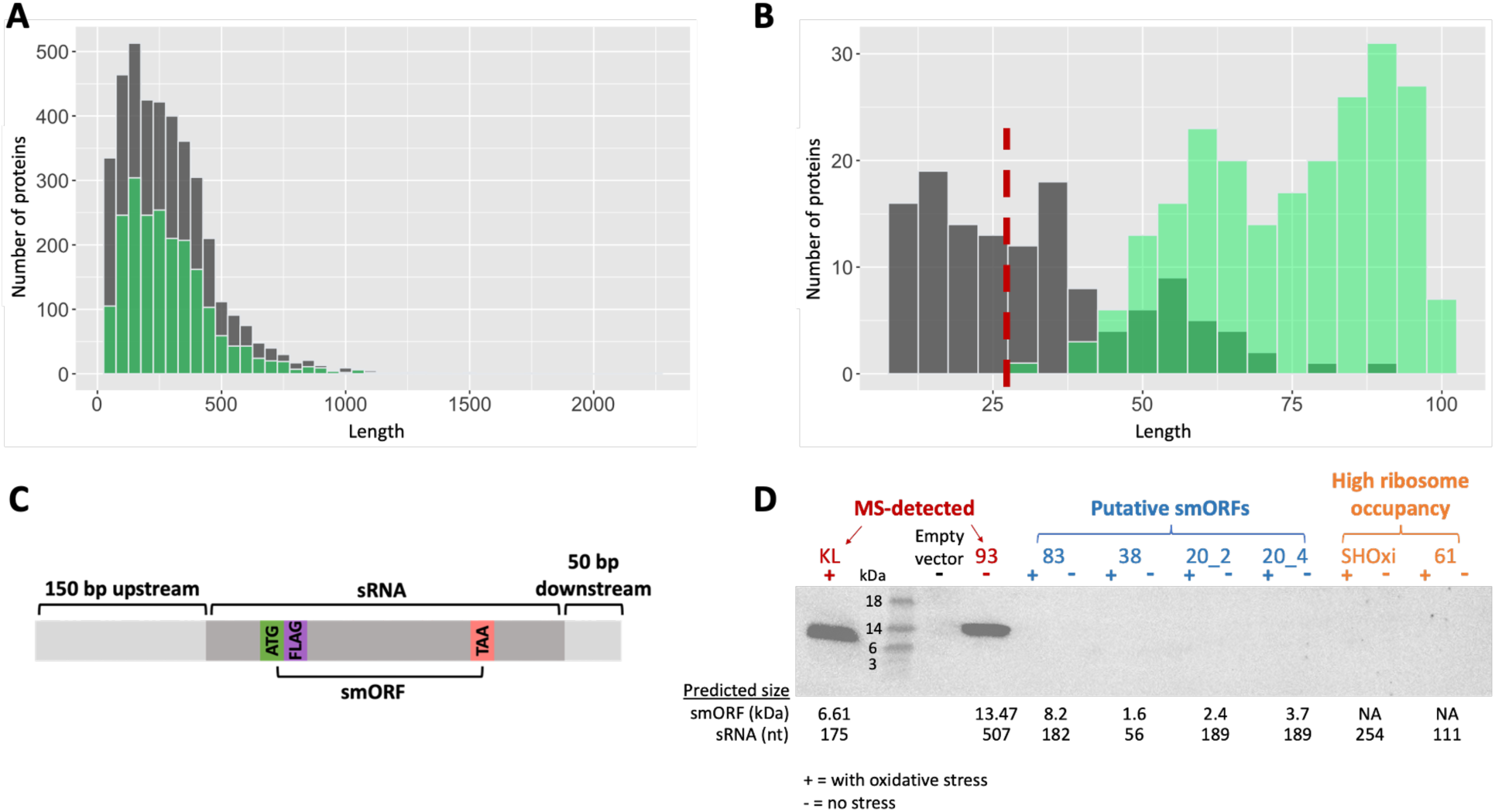
Discovery of novel small proteins encoded in sRNAs. A. Overlapping histogram of protein length (aa) for the *H. volcanii* annotated proteome (gray) and proteins detected by MS (overlaid in green). B. Overlapping histogram of protein length (aa) for putative peptides (predicted using NCBI ORF finder) encoded by sRNAs (gray) and small proteins detected by MS (overlaid in green). Detection cutoff of 30aa is indicated by red dashed line. C. Cloning strategy for tagging sRNAs for Western blot analysis. FLAG indicates the location of the 3xFLAG-tag. D. Western blot of tagged sRNAs. + indicates samples treated with hydrogen peroxide and - indicates control samples.

**Table 1.**
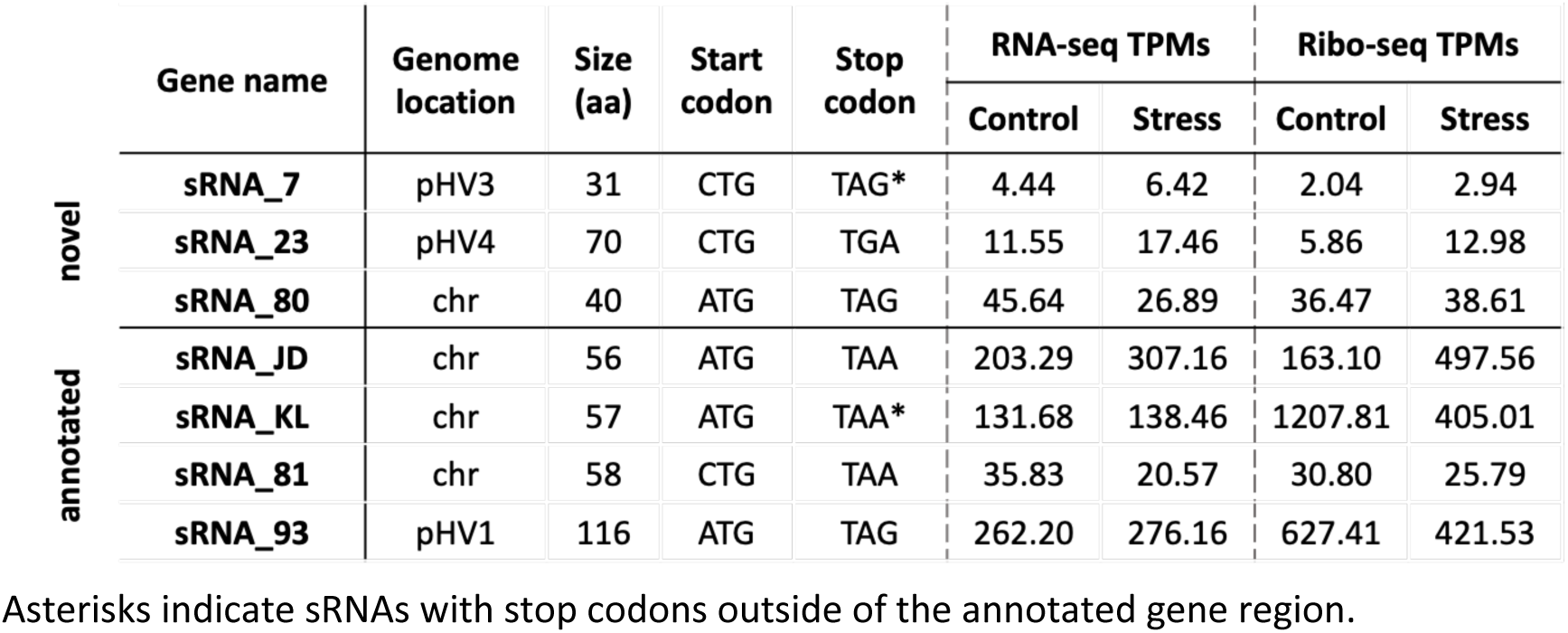
Annotated and novel small proteins encoded by sRNAs and detected by MS.

To further investigate sRNAs with ribosome occupancy, we selected a subset for validation by Western blot analysis (Table 2). This set included one sRNA with read counts below the minimum threshold for clustering (sRNA_38), one sRNA not present in the coding cluster (sRNA_83), three sRNAs within the coding cluster but not detected by MS (sRNA_20, sRNA_61, and SHOxi), and two smORFs detected by MS (sRNA_KL, sRNA_93). Each selected sRNA was tagged with a 3xFLAG epitope immediately downstream of the putative start codon and cloned into *H. volcanii* on an overexpression plasmid (Fig. 3C). To preserve native regulatory elements, 150 bp upstream and 50 bp downstream of each sRNA were included. For sRNA_20, which contained multiple putative ORFs, two start codons were tagged, including one corresponding to a previously annotated protein. Western blot analysis confirmed the expression of both MS-detected smORFs (sRNA_KL, sRNA_93), while no signal was observed for any of the other tagged sRNAs (Fig. 3D). Despite having both expression level and ribosome occupancy exceeding that of sRNA_KL and sRNA_93 (Table 2), no signal was detected for SHOxi and sRNA_61. This strongly suggests that SHOxi and sRNA_61 function as noncoding RNAs.

**Table 2.** sRNAs tagged for Western blotting.

|  | Gene name | Genome location | sRNA length (nt) | smORF size (aa) | Stop codon | RNA-seq TPMs |  | Ribo-seq TPMs |  |
| --- | --- | --- | --- | --- | --- | --- | --- | --- | --- |
|  |  |  |  |  |  | Control | Stress | Control | Stress |
| MS-detected | sRNA_KL | chr | 175 | 57 | yes | 131.68 | 138.46 | 1207.81 | 405.01 |
|  | sRNA_93 | pHV1 | 507 | 116 | yes | 262.20 | 276.16 | 627.41 | 421.53 |
| Putative smORFs | sRNA_38 | pHV4 | 56 | 16 | yes | 18.19 | 40.62 | 40.61 | 120.04 |
|  | sRNA_83 | chr | 182 | 55 | yes | 288.90 | 270.63 | 22.64 | 81.98 |
|  | sRNA_20_ORF2 | pHV4 | 189 | 21 | yes | 40.77 | 89.47 | 68.98 | 542.12 |
|  | sRNA_20_ORF4 | pHV4 | 189 | 34 | yes | 40.77 | 89.47 | 68.98 | 542.12 |
| High ribosome occupancy | sRNA_61 | chr | 111 | 32 | no | 4565.10 | 2281.40 | 1850.99 | 3381.31 |
|  | SHOxi | pHV3 | 254 | 73 | no | 346.33 | 1767.52 | 219.35 | 2034.76 |

sRNAs with “no” in the stop codon column had no stop codon contained in the predicted sRNA region as determined by ORFfinder.

Taken together, our combined LC-MS/MS and Western blot analyses identified seven smORFs encoded by sRNAs, including three previously unannotated smORFs. However, several sRNAs - most notably sRNA_61 and SHOxi - exhibited high ribosome occupancy without detectable protein products, highlighting the complexity of interpreting ribosome profiling data and suggesting potential regulatory or ribosome-associated noncoding functions.

### Ribosome-associated sRNAs may play regulatory roles during oxidative stress

We then focused on 17 stress-responsive sRNAs that had differential ribosome occupancy or TE under oxidative stress conditions (Fig. 4A, Table 3). Among these, five sRNAs had annotated smORFs, while the remaining 12 showed no evidence of coding potential. To further assess their coding potential, we applied the ribosome footprint length clustering analysis described earlier, this time using Ribo-seq data from oxidative stress-treated samples. Of the 12 unannotated sRNAs (with no evidence of coding potential), one failed to meet the minimum read count threshold and was excluded from the analysis. Among the remaining 11, three clustered with known coding genes, whereas nine did not (Fig. 4B). Given their lack of coding-like ribosome footprint distributions, we hypothesized that these nine stress-responsive sRNAs with high ribosome occupancy may play a regulatory role during oxidative stress. To investigate their potential functional roles, we used a suite of *in silico* approaches. Notably, two sRNAs were excluded from this analysis: SHOxi, which has been previously characterized as a regulatory RNA (35), and sRNA_41, which is annotated as encoding a small protein.

**Table 3.** sRNAs with differential ribosome occupancy or translational efficiency during oxidative stress.

| Name | RNA-seq |  | Ribo-seq |  | TE |  | TPMs |  |  |  | footprint distributions |  | protein detection |  |  |
| --- | --- | --- | --- | --- | --- | --- | --- | --- | --- | --- | --- | --- | --- | --- | --- |
|  | fold change | p-value | fold change | p-value | fold change | p-value | RNA-seq control | RNA-seq stress | Ribo-seq control | Ribo-seq stress | coding cluster control | coding cluster stress | annotated protein | ArcPP detection | MS detected |
| sRNA_20 | 1.21 | 0.00 | 2.63 | 0.00 | 1.41 | 0.04 | 40.22 | 88.01 | 68.74 | 537.73 | yes | yes | yes | no | no |
| sRNA_34 | -0.44 | 0.22 | -1.12 | 0.00 | -0.70 | 0.26 | 19.44 | 15.34 | 12.93 | 7.52 | yes | no | no | NA | no |
| sRNA_41 | -0.93 | 0.00 | -1.18 | 0.00 | -0.27 | 0.66 | 46.64 | 24.21 | 11.62 | 6.65 | no | no | yes | yes | no |
| sRNA_49 | 2.23 | 0.00 | 1.74 | 0.00 | -0.50 | 0.57 | 37.41 | 195.67 | 127.06 | 525.30 | no | no | no | NA | no |
| sRNA_58 | 0.40 | 0.37 | 1.23 | 0.01 | 0.81 | 0.39 | 21.45 | 34.76 | 41.31 | 135.96 | no | no | no | NA | no |
| sRNA_60 | -0.12 | 0.94 | 1.72 | 0.00 | 1.82 | 0.28 | 2491.07 | 1996.40 | 382.05 | 568.67 | NA | no | no | NA | no |
| sRNA_61 | -0.90 | 0.04 | 0.59 | 0.10 | 1.48 | 0.04 | 4496.30 | 2243.82 | 1843.17 | 3350.74 | yes | yes | no | NA | no |
| sRNA_62 | 2.24 | 0.00 | 3.15 | 0.00 | 0.89 | 0.42 | 17.24 | 79.53 | 4.06 | 45.72 | no | yes | no | NA | no |
| sRNA_71 | -0.31 | 0.38 | 1.35 | 0.05 | 1.63 | 0.04 | 76.87 | 105.29 | 8.57 | 27.23 | NA | NA | no | NA | no |
| sRNA_77 | 0.90 | 0.04 | 2.57 | 0.00 | 1.66 | 0.10 | 23.84 | 43.05 | 3.76 | 13.07 | NA | no | no | NA | no |
| sRNA_83 | 0.06 | 0.89 | 1.59 | 0.00 | 1.52 | 0.03 | 284.91 | 266.25 | 22.57 | 81.26 | no | no | no | NA | no |
| sRNA_95 | 1.14 | 0.00 | 1.61 | 0.00 | 0.46 | 0.56 | 21.16 | 45.04 | 9.43 | 33.82 | no | no | no | NA | no |
| sRNA_96 | 1.65 | 0.00 | 1.59 | 0.00 | -0.08 | 0.92 | 51.26 | 140.20 | 5.59 | 19.10 | yes | yes | no | NA | no |
| sRNA_CD | 0.95 | 0.01 | 1.50 | 0.00 | 0.53 | 0.44 | 83.53 | 141.43 | 168.64 | 569.29 | no | yes | yes | yes | no |
| sRNA_JD | 0.68 | 0.20 | 1.33 | 0.01 | 0.64 | 0.57 | 200.14 | 301.88 | 162.34 | 493.83 | yes | yes | yes | yes | yes |
| sRNA_KL | 0.16 | 0.57 | -1.93 | 0.00 | -2.12 | 0.00 | 129.94 | 136.19 | 1203.27 | 401.61 | yes | yes | yes | yes | yes |
| sRNA_SHOxi | 2.46 | 0.00 | 2.89 | 0.00 | 0.42 | 0.72 | 341.52 | 1738.88 | 218.70 | 2017.74 | yes | no | no | NA | no |

Fold change refers to log<sub>2</sub> fold change, and p-value refers to the adjusted p-value.

**Figure 4.**
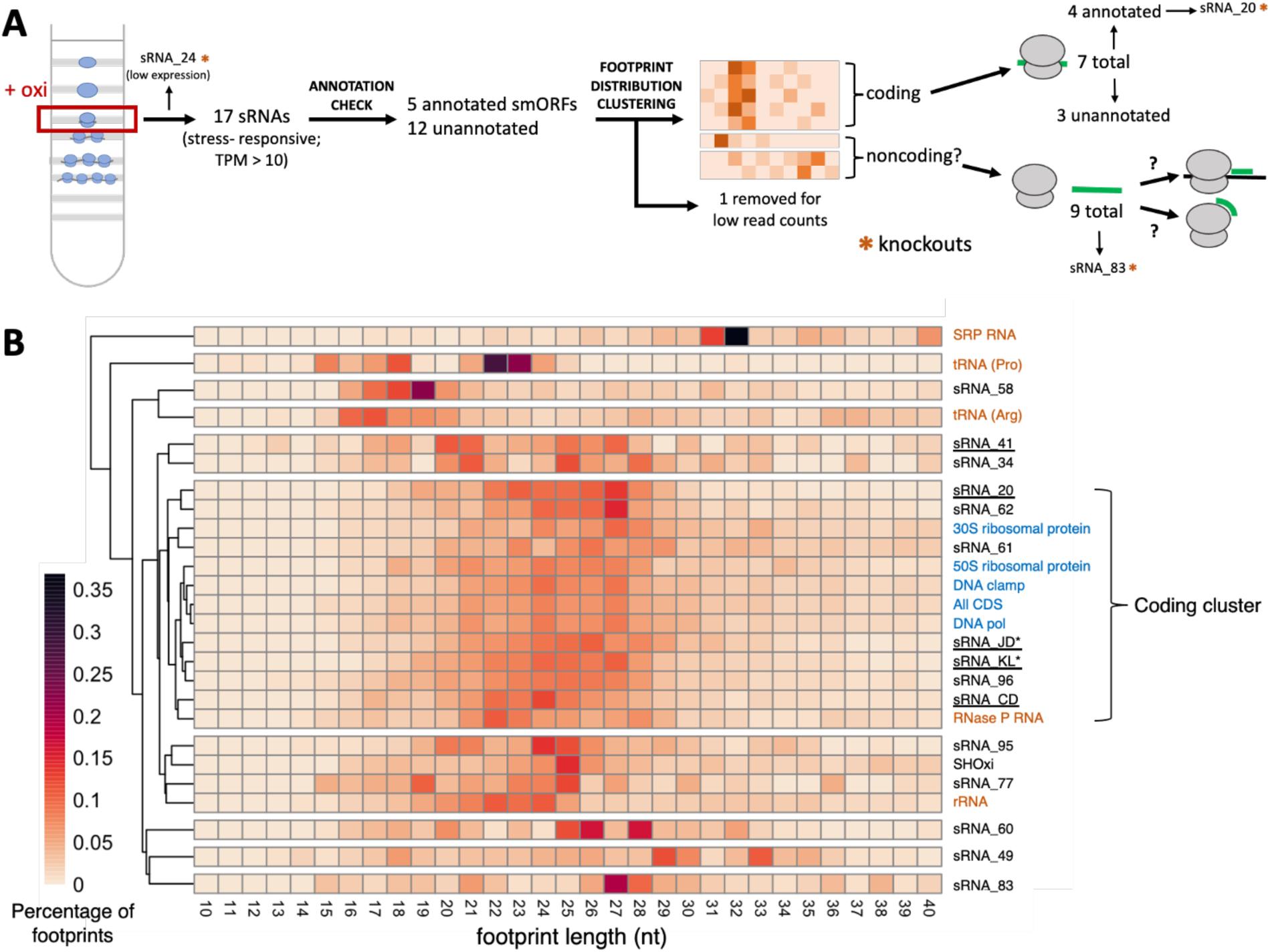
Ribosome footprint length distributions and stress-responsive sRNAs with ribosome occupancy. A. Schematic of analyses to assess the coding potential for sRNAs with ribosome occupancy, starting with the isolation of monosome fractions on a sucrose gradient using samples treated with oxidative stress. B. Clustered heatmaps of ribosome footprint length distributions for sRNAs, coding genes, and noncoding genes with ribosome occupancy under oxidative stress. Darker colors indicate higher read percentage. Known coding genes are shown in blue, known noncoding genes in orange, and sRNAs in black. sRNAs with annotated proteins are underlined. sRNAs for which peptides have been detected by MS are marked with an asterisk. Percentages for ‘All CDS’ were calculated using sum read counts of all coding genes.

We first performed BLASTn to assess the conservation of these noncoding stress-responsive sRNAs. Five had homology to one or more halophilic species. However, for two of these, the majority of homologous hits overlapped with an upstream gene on the opposite strand, complicating the interpretation. The remaining two sRNAs had no detectable homology to sequences in other species. We selected three sRNAs - sRNA_58, sRNA_77, and sRNA_83 - for further analysis based on their conservation in at least one other species and the absence of overlap with annotated genes.

We performed a multiple sequence alignment and predicted consensus secondary structures for sRNA_58, sRNA_77, and sRNA_83 using LocARNA (68), incorporating all homologous sequences identified in our BLASTn search. Each sRNA exhibited strong secondary structure potential and high sequence conservation across species (Fig. 5A; Fig. S7), suggesting functional relevance. To explore their regulatory role, we predicted putative mRNA targets using IntaRNA [Fig. 5B; Tables S14-16; (69)] and determined which of those predicted targets had differential ribosome occupancy or TE under oxidative stress (Fig. 5C, Table 4). sRNA_58 had the fewest high-confidence predicted targets (p <0.01), with only five out of 29 (17%) showing differential ribosome occupancy or TE in response to oxidative stress. In contrast, sRNA_77 and sRNA_83 had a larger number of predicted high-confidence targets. Specifically, 13 out of 44 (30%) putative mRNA targets for sRNA_83, and 15 out of 42 (36%) targets for sRNA_77 were differentially translated in response to oxidative stress.

**Figure 5.**
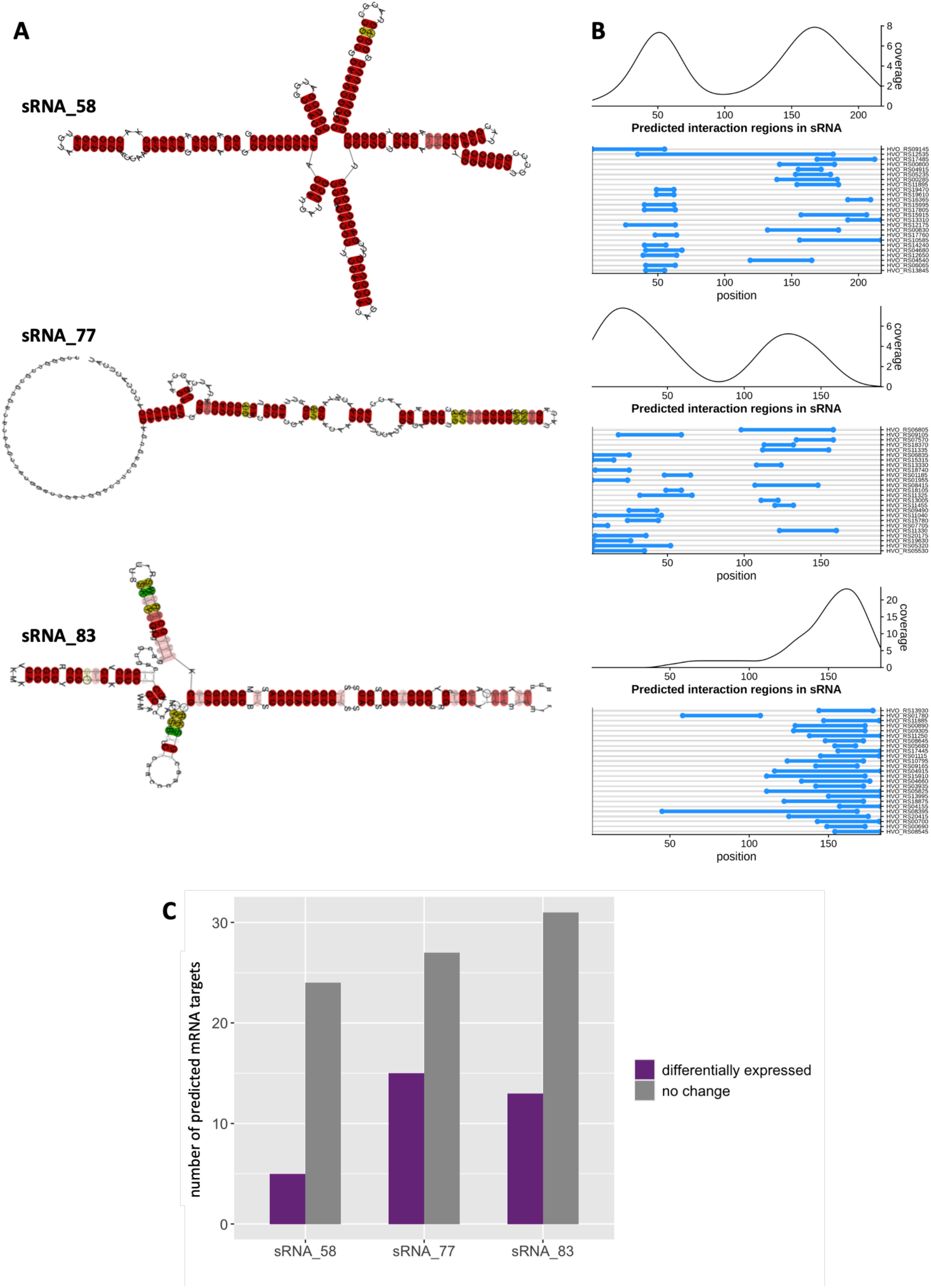
Characterization of stress-responsive sRNAs with ribosome occupancy. A. Consensus RNA structural predictions for sRNAs with no evidence of coding potential were generated using LocARNA. Colored bases indicate sequence and structural conservation of compatible base pairs within the predicted structure (see Fig. S7). Colors correspond to number (red: 1, yellow: 2, green: 3) of base pair types (C-G, G-C, A-U, U-A, G-U, or U-G) with red being the most highly conserved and green the least conserved. Color saturation decreases with the number of incompatible base pairs. B. IntaRNA was used to predict putative mRNA targets (Tables S14-16). Blue lines indicate putative binding regions along the sRNA. Curves depict coverage of putative binding regions along the sRNA. C. Bar chart of predicted mRNA targets (p <0.01) with differential ribosome occupancy or TE during oxidative stress (purple). The number of predicted targets that are unaffected by oxidative stress are shown in gray.

**Table 4.** Putative mRNA targets with differential ribosome occupancy or TE during oxidative stress.

|  | Target Name | Function | RNA-seq |  | Ribo-seq |  | TE |  | TPMs |  |  |  |
| --- | --- | --- | --- | --- | --- | --- | --- | --- | --- | --- | --- | --- |
|  |  |  | Fold Change | p-value | Fold Change | p-value | Fold Change | p-value | RNA-seq Control | RNA-seq Stress | Ribo-seq Control | Ribo-seq Stress |
| sRNA_58 | HVO_RS04540 | hypothetical protein | -1.21 | 0.00 | -1.21 | 0.00 | -0.02 | 0.98 | 326.20 | 169.80 | 370.97 | 208.87 |
|  | HVO_RS13165 | ABC transporter permease | -1.64 | 0.00 | -1.27 | 0.00 | 0.36 | 0.51 | 640.15 | 188.77 | 85.81 | 45.53 |
|  | HVO_RS17485 | D-2-hydroxyacid dehydrogenase | -1.01 | 0.00 | -1.60 | 0.00 | -0.61 | 0.07 | 27.50 | 47.10 | 58.33 | 27.88 |
|  | HVO_RS17805 | hypothetical protein | 1.00 | 0.00 | 1.46 | 0.00 | 0.45 | 0.45 | 54.93 | 91.69 | 36.61 | 98.47 |
|  | HVO_RS19470 | hypothetical protein | 1.33 | 0.01 | -0.65 | 0.05 | -2.01 | 0.01 | 7.57 | 15.74 | 55.04 | 44.71 |
| sRNA_77 | HVO_RS05320 | YccF domain-containing protein | 1.01 | 0.00 | 1.37 | 0.00 | 0.34 | 0.62 | 26.87 | 53.00 | 37.93 | 121.84 |
|  | HVO_RS06805 | co-chaperone YbbN | 0.62 | 0.06 | -0.66 | 0.03 | -1.30 | 0.01 | 1088.48 | 1614.45 | 2937.95 | 2326.41 |
|  | HVO_RS07705 | hypothetical protein | -0.52 | 0.09 | 0.80 | 0.07 | 1.31 | 0.04 | 51.84 | 33.24 | 11.22 | 23.01 |
|  | HVO_RS08415 | amidohydrolase family protein | -0.88 | 0.00 | -1.87 | 0.00 | -1.01 | 0.03 | 82.41 | 82.17 | 87.53 | 55.80 |
|  | HVO_RS09105 | 2Fe-2S domain-containing protein | -0.24 | 0.31 | -1.24 | 0.00 | -1.01 | 0.05 | 67.63 | 53.64 | 76.35 | 40.75 |
|  | HVO_RS11380 | hypothetical protein | 1.06 | 0.00 | 2.71 | 0.00 | 1.63 | 0.04 | 21.14 | 49.53 | 68.11 | 579.99 |
|  | HVO_RS11455 | BMP family protein | -0.72 | 0.00 | -1.78 | 0.00 | -1.08 | 0.05 | 1322.62 | 720.75 | 1618.05 | 586.70 |
|  | HVO_RS13145 | hypothetical protein | 0.93 | 0.00 | 1.06 | 0.00 | 0.11 | 0.86 | 193.21 | 286.04 | 47.28 | 123.46 |
|  | HVO_RS14580 | dehydrorhamnose epimerase family protein | -0.76 | 0.03 | -1.88 | 0.00 | -1.14 | 0.07 | 1726.87 | 885.36 | 986.87 | 329.83 |
|  | HVO_RS15315 | hypothetical protein | -0.73 | 0.00 | 0.74 | 0.03 | 1.46 | 0.00 | 133.96 | 75.10 | 13.40 | 25.06 |
|  | HVO_RS17930 | 30S ribosomal protein S28e | -1.01 | 0.00 | -1.63 | 0.00 | -0.63 | 0.13 | 8234.66 | 3814.46 | 1902.49 | 770.44 |
|  | HVO_RS18350 | hypothetical protein | 1.10 | 0.00 | 1.19 | 0.02 | 0.07 | 0.94 | 19.56 | 38.77 | 6.65 | 17.79 |
|  | HVO_RS18595 | GNAT family N-acetyltransferase | 0.27 | 0.39 | 1.05 | 0.00 | 0.76 | 0.12 | 30.92 | 27.85 | 9.31 | 23.79 |
|  | HVO_RS18740 | hypothetical protein | 0.16 | 0.66 | 1.18 | 0.00 | 0.99 | 0.08 | 68.49 | 68.32 | 17.95 | 42.57 |
|  | HVO_RS20570 | rubrerythrin-like domain-containing protein | 0.97 | 0.00 | 3.78 | 0.00 | 2.79 | 0.00 | 50.42 | 49.15 | 17.52 | 161.98 |
| sRNA_83 | HVO_RS00230 | methyltransferase | -1.08 | 0.00 | -1.08 | 0.00 | -0.01 | 0.98 | 219.52 | 94.40 | 163.37 | 97.06 |
|  | HVO_RS00700 | ABC transporter substrate-binding protein | -2.13 | 0.00 | -2.70 | 0.00 | -0.59 | 0.42 | 190.15 | 38.32 | 819.59 | 157.72 |
|  | HVO_RS01670 | racemase enzyme family protein | 1.66 | 0.00 | 1.20 | 0.01 | -0.47 | 0.60 | 4.57 | 12.69 | 6.18 | 18.40 |
|  | HVO_RS03935 | ring-cleaving dioxygenase | 0.98 | 0.00 | 1.44 | 0.00 | 0.44 | 0.36 | 36.14 | 63.65 | 24.10 | 83.36 |
|  | HVO_RS04155 | hypothetical protein | 0.96 | 0.08 | 2.51 | 0.00 | 1.55 | 0.09 | 36.47 | 27.77 | 18.03 | 36.31 |
|  | HVO_RS04735 | hypothetical protein | -0.12 | 0.72 | 1.10 | 0.00 | 1.21 | 0.01 | 50.36 | 45.95 | 18.86 | 51.67 |
|  | HVO_RS05345 | thermosome subunit alpha | -1.06 | 0.00 | -1.58 | 0.00 | -0.53 | 0.37 | 4645.21 | 2047.07 | 6582.65 | 2681.76 |
|  | HVO_RS08545 | SPFH domain-containing protein | 0.46 | 0.04 | 1.25 | 0.00 | 0.77 | 0.10 | 295.03 | 436.26 | 65.36 | 202.53 |
|  | HVO_RS08645 | NAD(P)-dependent dehydrogenase | -0.74 | 0.00 | -1.63 | 0.00 | -0.91 | 0.01 | 143.06 | 78.46 | 197.19 | 80.87 |
|  | HVO_RS13905 | serine-tRNA ligase | -0.51 | 0.02 | -1.27 | 0.00 | -0.77 | 0.07 | 384.53 | 246.71 | 547.21 | 295.37 |
|  | HVO_RS13930 | SIMPL domain-containing protein | 0.99 | 0.00 | 1.69 | 0.00 | 0.68 | 0.17 | 65.12 | 120.33 | 24.14 | 91.85 |
|  | HVO_RS17570 | HpcH/Hpal aldolase/citrate lyase family protein | 0.96 | 0.01 | 1.87 | 0.00 | 0.89 | 0.38 | 133.65 | 232.04 | 49.88 | 213.11 |
|  | HVO_RS19230 | ABC transporter ATP-binding protein | -1.35 | 0.00 | -1.65 | 0.00 | -0.32 | 0.55 | 38.24 | 18.34 | 66.16 | 27.28 |

Fold change refers to log<sub>2</sub> fold change, and p-value refers to the adjusted p-value.

Collectively, of the eight sRNAs with differential ribosome occupancy or TE under oxidative stress [excluding SHOxi, previously characterized (35)], five were conserved among halophilic species. Two of these, sRNA_77 and sRNA_83, had numerous putative targets that exhibited translational regulation during oxidative stress, suggesting they may play regulatory roles in the stress response.

### sRNA knockouts have a similar phenotype to wild-type during oxidative stress

To explore the functional role of ribosome-associated sRNAs during oxidative stress, we generated sRNA knockout strains for three sRNAs with a range of ribosome occupancy. sRNA_24 represented low ribosome occupancy (TPM <10), sRNA_83 represented moderate occupancy (50-100 TPM), and sRNA_20 represented high occupancy (>200 TPM). Knockouts were generated as previously described (70) and whole genome sequencing confirmed the complete deletion of the target sRNAs. Under oxidative stress conditions induced by H₂O₂ treatment, none of the knockout strains exhibited observable phenotypic differences compared to the wild-type (WT) strain (Fig. S8A, B).

To identify putative regulatory targets for these sRNAs, we performed RNA-seq using three biological replicates for each knockout strain and the WT strain exposed to oxidative stress, along with non-stress WT controls, as previously described (35). The RNA-seq data were analyzed, as described above, with significance cutoffs of adjusted p-value <0.05 and absolute log_2_ fold change >1. No significantly differentially expressed genes were detected for sRNA_83 and sRNA_20 knockout strains when compared to the WT strain under oxidative stress (Fig. S8C, D). However, we found 7 genes significantly upregulated for the sRNA_24 knockout strain (Fig. S8E), which was further validated by qPCR (Fig. S8F). Three of these targets were members of an operon involved in glycerol metabolism in *H. volcanii* [Table S17, (71)]. Remarkably, genome analysis revealed that nine genes differentially regulated and with p-value <0.05 were located immediately downstream of sRNA_24 locus (Fig. S8G, Table S17). None of these genes were predicted as putative sRNA targets using IntaRNA, indicating a likely polar effect of the sRNA gene deletion.

## Discussion

This study provides a comprehensive view of the translational landscape of the model archaeon *H. volcanii* during oxidative stress by integrating RNA-seq and Ribo-seq analyses. While previous studies in *H. volcanii* have reported changes in protein abundance under oxidative stress using MS (46), such approaches cannot distinguish between transcriptional and translational regulation. Furthermore, it is well established that transcript levels measured by RNA-seq often correlate only modestly - and with high variability - with protein level determined by MS (72, 73), highlighting the complexity of processes that govern protein synthesis. To better understand how gene expression is controlled at the level of translation, we used RNA-seq and Ribo-seq to examine both ribosome occupancy and TE across the transcriptome.

Our analysis revealed that hundreds of genes exhibited differential TE in *H. volcanii* during oxidative stress. Notably, we observed global downregulation of translation, particularly affecting ribosomal proteins, suggesting a substantial reduction in *de novo* protein synthesis under stress conditions. This translational repression is consistent with previous studies in *H. volcanii* and *E. coli* under oxidative stress (27, 74, 75), as well as in *Halobacterium* NRC-1 following UV irradiation (76), all of which reported downregulation of translation-related genes. This downregulation is thought to serve as a protective and energy-conserving strategy, redirecting cellular resources toward damage repair and stress defense while minimizing oxidative damage to the translation machinery (77). Similarly, in eukaryotic systems, stress-induced translation arrest is well documented and often involves the formation of stress granules containing translationally stalled complexes (40, 78, 79). Interestingly, a previous mass spectrometry study in *H. volcanii* reported upregulation of several translation-related proteins, including ribosomal proteins, in cells exposed to hypochlorite stress (46), highlighting potential discrepancies between transcriptional, translational, and proteomic responses. However, a study with *Halobacterium* NRC1 subjected to ionizing radiation (IR) addressed the sequential nature of the cell’s transcriptional (RNA-seq) and translational (proteomic) responses (80). The study found a high level of correlation between mRNA and protein levels by temporally shifting the protein level changes by 20 to 30 min, demonstrating the importance of the time dimension.

In addition to translation-related genes, we observed downregulation of genes encoding peroxidases and components of the tricarboxylic acid (TCA) cycle - findings that are consistent with responses observed in *Deinococcus radiodurans* and *Halobacterium* NRC-1 following IR (80–82). Because IR generates massive oxidative stress, suppression of general metabolism and redox-active pathways may help cells limit further damage and maintain homeostasis. Taken together, the divergence between RNA-seq, Ribo-seq, and MS datasets suggests that multiple layers of regulation are employed to control gene expression during oxidative stress. Such complex gene regulation has been documented in both eukaryotes and bacteria. For example, cold shock protein in mammals (83), quorum sensing signals in *Agrobacterium tumefaciens* (84, 85), and the tryptophan operon in *E. coli* (86, 87) are all regulated at both the transcriptional and translational levels. Similarly, in *H. volcanii*, the integration of transcriptional and translational control likely enables a rapid and precise response to oxidative stress, allowing selective synthesis of essential proteins while conserving energy by broadly reducing translation.

Remarkably, we found that many sRNAs previously classified as noncoding (27) exhibited high ribosome occupancy, suggesting that ribosome association may not be limited to actively translated transcripts. Leveraging our Ribo-seq data, we analyzed ribosome footprint length distribution and found that several of these sRNAs had footprint length distributions similar to those of coding genes. A subset of these sRNAs were found to encode annotated smORFs, and using MS, we identified seven small proteins encoded by sRNAs. Unfortunately, sequence or structural homology searches did not provide insights into the potential functions of these sRNA-encoded proteins. One exception is the small protein encoded in sRNA_KL, which has previously been detected by both Ribo-seq and MS (53). This small protein contains a zinc finger domain, and a deletion mutant showed increased growth on glycerol (88). It is important to note that the ability to encode small proteins does not exclude these sRNAs from having post-transcriptional regulatory roles. In bacteria, dual function sRNAs that both encode proteins and act as regulatory sRNAs are well documented (89–91). Similar dual-function sRNAs have been proposed in archaea, particularly under nitrogen starvation conditions (92). These findings suggest that, as in other organisms (89, 93), *H. volcanii* may also use dual-function sRNAs, especially in response to oxidative stress.

Ribo-seq is potentially a more sensitive method than MS for detecting smORFs (53, 92). However, there is enduring concern that it may overestimate the number of translated sRNAs (94, 95), making MS-based proteomics an important complementary approach to provide direct protein-level evidence. In addition, analyzing ribosome footprint length distribution offers another strategy to assess the coding potential of sRNAs. As discussed above, this analysis successfully distinguished most coding genes and sRNAs with annotated smORFs from noncoding genes. Nonetheless, this method was not accurate for all coding genes and requires high read counts to reliably separate coding and noncoding transcripts.

We also identified several sRNAs with high ribosome occupancy that had no evidence of coding potential. These sRNAs did not yield detectable protein even when putative ORFs were individually tagged and probed by Western blotting. In contrast, smORFs detected by LC-MS were also confirmed by Western blot, despite having lower ribosome occupancy. This discrepancy may reflect the limitations in detecting low-abundance small proteins by MS or Western blotting. For lowly expressed sRNAs, overexpression with an inducible promoter, as previously shown (53), may be necessary for detection. Alternatively, sRNAs with high ribosome occupancy and no evidence of coding potential may be functionally associated with ribosomes, possibly directly interacting with them. Their association is unlikely to be an artefact of the ribosome profiling protocol; although RNAs with secondary structure are generally resistant to nuclease digestion, the small size of these sRNAs would prevent their co-sedimentation with monosomes in a sucrose gradient. This suggests a specific association with ribosomes or ribosome-bound mRNAs. Prior studies have reported the presence of sRNAs associated with ribosomes in *H. volcanii* under stress conditions (96, 97). Sequencing and northern blotting of sRNAs that co-purify with ribosomes identified a tRNA-derived fragment that binds the small ribosome subunit, inhibiting peptide bond formation (96), as well as an intergenic sRNA that interacts with the large subunit to regulate the translation of a specific mRNA (97).

Several ribosome-associated sRNAs lacking coding potential were also responsive to oxidative stress, suggesting potential regulatory functions. These included SHOxi, which binds to and destabilizes malic enzyme transcripts, enhancing cell survival during oxidative stress (35), thus arguing against this sRNA being a nonfunctional *de novo* gene intermediate. Importantly, our RNA-seq and Ribo-seq experiments were performed with four biological replicates each, and the ribosome-associated sRNAs discussed here consistently showed high ribosome occupancies. Deletion of three such sRNAs did not result in a growth phenotype under oxidative stress and caused little to no change in the expression level of other genes. This suggests that these sRNAs may not modulate transcript stability directly, unlike some well-characterized sRNAs in both bacteria (32, 98, 99) and archaea (100, 101). For example, in *H. volcanii*, SHOxi destabilized its mRNA target through complementary base pairing (35). However, as observed in bacterial systems (102, 103), deletion of a single sRNA may be insufficient to observe its regulatory function. Targeted knockout of multiple stress-responsive, ribosome-associated sRNAs may therefore be necessary to fully elucidate their role during oxidative stress. Taken together, our findings suggest that several oxidative stress-associated sRNAs in *H. volcanii* are closely associated with ribosomes but do not affect mRNA stability. We hypothesize they regulate translation directly through interaction with the ribosome or ribosome-bound mRNAs.

Using RNA-seq and Ribo-seq in a model archaeon, we demonstrated global and targeted translation regulation in response to oxidative stress. We also identified a class of untranslated, ribosome-associated sRNAs likely involved in translational regulation under oxidative stress. While the precise mechanisms of action remain to be elucidated, this study highlights the complexity of translational regulation in archaea and underscore the multifaceted architecture of stress-responsive regulatory networks.

## Materials and Methods

### Cell culturing and oxidative stress treatment

*Haloferax volcanii* DS2 strain H98 (70) was grown in liquid Hv-YPC (70) supplemented with 40 mg/mL thymidine at 42°C, with shaking, to OD >0.5. Cultures were diluted into fresh media to an OD of 0.01 and grown to OD 0.4-0.6. For oxidative stress treatment, cultures were incubated with 2mM H_2_O_2_ for one hour at 42°C with shaking, as previously described (27).

### Ribosome profiling

Ribo-seq was performed as previously described (52, 104). In brief, cultures were flash frozen in liquid nitrogen and lysed by cryogenic mill, using 55 g of frozen culture and 100 µg/mL of anisomycin. Ribosomes were collected by ultracentrifugation on a 1 M sucrose cushion using a Ti45 rotor at 185,511x*g* (40,000 rpm) for three hours at 4°C. Polysomes were digested using MNase and separated using a 10-50% sucrose gradient (Fig. S9A). RNA from monosome fractions was isolated with Trizol LS reagent, followed by size selection (Fig. S9B), and generation of libraries for high-throughput sequencing (Fig. S9C, D). Libraries were sequenced by Novogene (150 bp, paired-end reads, NovaSeq).

### RNA-seq

RNA-seq libraries were prepared using 20 mL of culture pelleted by centrifugation. RNA extraction was performed as previously described (27) using a Quick-RNA Miniprep kit (Zymo). rRNA was depleted using 2.5 µg of DNase-treated RNA and the Ribo-Zero rRNA removal kit for bacteria (Illumina). The SMARTer Stranded RNA-Seq kit (Takara Bio) was used to prepare the libraries with 100 ng RNA, dual-index primers, and 12 PCR cycles. Libraries were sequenced by Novogene (150 bp, paired-end reads, NovaSeq). For knockout strains, RNA was prepared as described above, using 3 replicates per strain. All strains were treated with H_2_O_2_, with 3 replicates of untreated WT as controls. Library preparation and sequencing were performed by SeqCenter in Pittsburgh, PA (150 bp, paired-end reads, NovaSeq X plus) using Illumina’s Stranded Total RNA Prep Ligation with Ribo-Zero Plus kit and custom probes.

### Analysis of sequencing data

Demultiplexed reads provided by Novogene and SeqCenter were analyzed using only forward reads for Ribo-seq libraries and both reads for RNA-seq libraries. For full analysis details, see Table S18. Adapters were trimmed using Trim Galore [(105), https://github.com/FelixKrueger/TrimGalore] and reads were aligned to rRNA genes using Bowtie (106). Unaligned reads (those not aligning to rRNA) were written to a new file and used for genome alignment, which was performed using Bowtie and the *H. volcanii* DS2 genome assembly ASM2568v1 from NCBI (49). Samtools (107) was used to generate bam and sorted bam files. Coverage and depth were calculated using samtools and bedtools. Average depth per chromosome was calculated using a custom Python script. TPM were calculated using StringTie (108), with a provided reference annotation file (gff from NCBI Refseq assembly GCF_000025685.1 downloaded March 2023 with added entries for sRNAs). Read counts were calculated using featureCounts (109). See Tables S19, S20, and S21 for statistical summaries of all analyses.

### Differential expression and TE analysis

Differential expression was determined using DESeq2 (110). For RNA-seq and Ribo-seq data, the variable of interest was experimental condition. For TE, the ratio between Ribo-seq and RNA-seq was used, in addition to two variables of interest: sample (RNA-seq or Ribo-seq) and experimental condition. For full details, see Table S18.

### Characterization of significant genes

The number of leaderless genes was determined using published data that characterized 1851 annotated genes as leadered or leaderless (111). 68% of genes with differential TE were uncharacterized and therefore not considered in calculations for the percent of leaderless genes. Custom Python scripts were used to calculate percent of leaderless genes, codon frequencies, and amino acid frequencies. Heatmaps were generated using R. The STRING database [web version 12.0; (58)] was used to predict interactions and to determine KEGG enrichments with default settings and MCL clustering. Gene interaction predictions are based on evidence including genomic proximity, experimental data and co-expression of homologs, association in curated databases, and text mining of abstracts from published literature. 84% of genes with differential TE were successfully mapped to the STRING database. The PANTHER database (web version 19.0) was used to generate a generic mapping file (57, 112) for the *H. volcanii* DS2 protein fasta file downloaded from NCBI. Overrepresentation tests were conducted using Fisher’s test and FDR correction.

### smORF prediction

For ORF prediction using the web version of Open Reading Frame Finder [ORFfinder, NCBI; (66)], a minimal length of 30 nt and alternative initiation codons were allowed. For ORF prediction using Ribo-seq data, an adapted version of REPARATION (65), using blast instead of USEARCH (https://github.com/RickGelhausen/REPARATION_blast) was used. For option details, see Table S18. A custom python script compared and combined predictions with and without Shine-Dalgarno. Predicted genes were visualized using IGV, and a manual scan was conducted to compare overlap between replicates for control or oxidative stress treatment. Predicted genes were required to be present in at least 3 biological replicates. Novel gene lists for control and stress samples were then combined.

### Footprint distribution clustering

Lengths of reads aligning to individual genes were obtained using a custom Bash script, followed by analysis using R to generate heatmaps. Clustering was performed using the pheatmap (https://github.com/raivokolde/pheatmap) package with default settings and the minimum number of clusters required to separate rRNA from known coding genes in control samples.

### Mass spectrometry sample preparation

Samples for mass spectrometry were prepared using 600 mL cultures from three replicates; half of the culture was used for control samples and the other half was treated with H_2_O_2_. Cells were pelleted by centrifugation then resuspended in 8 mL lysis buffer (2 M NaCl, 8 M urea, 0.1 M Tris-Cl, 10 mM DTT) and flash frozen. Resuspension was ground in a cryogenic mill (SPEX 6870) with a 1 minute pre-cool, then 8 cycles of alternating 1 minute of run time and 1 minute of cool time, at a rate of 8 cycles/second. The thawed lysate was spun twice at 15000 x*g* for 15 minutes at 4°C. Lysis buffer was added to a volume of 10 mL, then filtered through a 0.2-micron syringe filter and brought to a final volume of 12 mL with lysis buffer. Size selection was performed by centrifugation in 100K MWCO filters (Pall) at 5000 x*g* for 2 hours at room temperature, then centrifugation in 30K MWCO filters (Pall) at 5000 x*g* for 45 minutes at room temperature. Iodoacetamide was added to a final concentration of 54 mM and incubated for 45 minutes in the dark at room temperature, followed by addition of 11.1% of volume of trichloroacetic acid. Samples were flash frozen, then incubated at 4°C overnight. Samples were centrifuged at 20000 x*g* for 30 minutes at 4°C and supernatant was removed. Pellets were washed twice with acetone, air-dried for 10 minutes, then resuspended in 50 µL of 8 M urea in phosphate-buffered saline (PBS), pH 7.4 and quantified using the Qubit Protein Assay kit. 30 µg of total protein in 200 µL of 2 M urea in PBS, pH 7.4 were digested with trypsin and prepared for MS as previously described (113).

### Mass spectrometry acquisition

Chromatographic separation of protein digests was carried out as previously described (114), injecting 1 μg of protein per sample.

### Mass spectrometry data analysis

The Proteome Discoverer (PD) Software Suite v2.4, Minora Algorithm (ThermoFisher) was used to process mass spectra and perform Label Free Quantification (LFQ), comparing control and stress samples with default settings for all analysis nodes except where specified. Data were searched against the *H. volcanii* DS2 proteome fasta file from Uniprot (entry UP000008243, organism ID 309800; downloaded January 2023). The reference fasta file was appended with additional sequences for putative smORFs. For peptide identification, the Proteome Discoverer Sequest HT node was selected for a tryptic search allowing up to 2 missed cleavages. A precursor mass tolerance of 10 ppm was used for the MS1 level, and a fragment ion tolerance of 0.02 was set at the MS2 level. Methionine oxidation and N-terminus acetylation were allowed as dynamic modifications, with carbamidomethylation on cysteines defined as a static modification. Protein abundance ratio data were exported using a three-level hierarchy (protein > peptide group > consensus features). A minimum of 2 PSMs and maximum FDR of 5% were required to determine detection of novel small proteins. See Table S22 for statistical summaries.

### Generating tagged sRNAs

sRNA regions (including at minimum 150 bp upstream and 50 bp downstream) were amplified from genomic DNA and cloned into pTA1300 using *Kas*I and *Bam*HI. Ligation products were transformed into DH5α competent cells (NEB), followed by plasmid extraction (QIAprep spin miniprep kit) and verification by Sanger sequencing (GENEWIZ, Azenta). A 3xFLAG tag was added after the predicted start codon using Q5 site-directed mutagenesis (NEB). Purified plasmids were transformed into *H. volcanii* as previously described (115), using the lab strain H98.

### Western blots

25 mL of culture were pelleted by centrifugation at 10,000 x*g* for 6 minutes at 4°C. Pellets were resuspended in 1 mL lysis buffer (50 mM Tris-HCl pH 7.5, 2 M urea, 0.2% SDS, 0.1 mg/mL EDTA-free protease inhibitor cocktail (Roche)) and lysed with sterile glass beads (116). Protein concentration was quantified using the Qubit Protein Assay kit. Proteins (10-100 µg total) were separated on a 4-15% stain free acrylamide gel (BioRad), transferred to a nitrocellulose membrane, and incubated with anti-Flag monoclonal antibodies conjugated with alkaline phosphatase (1:10,000 dilution; Sigma cat no. A9469), as previously described (116). The antibody:protein complexes were visualized using 0.25 mM CDP-Star Substrate (Tropix) with 1x Nitro-Block-II Enhancer (Invitrogen) and the BioRad ChemiDoc Imager.

### sRNA in silico tools

Web versions of nucleotide BLAST (117), IntaRNA (69, 118), and LocARNA (68) were used with default settings.

### Generating knockouts

Knockout strains were generated using the established protocol for *H. volcanii* (70). For sRNA_83, a knockout plasmid was constructed by amplifying upstream and downstream flanking regions from genomic DNA and cloning into pTA131 using *Hind*III, *Xba*I, and *Eco*RI. Knockout plasmids for sRNA_20 and sRNA_24 were generated by Azenta, with upstream and downstream flanking regions cloned into pTA131 using *Hind*III and *Xba*I. Knockout plasmids were transformed into *H. volcanii* (115), strain H98, followed by selection for pop-in (uracil selection) and pop-out (5-FOA selection). Whole genome sequencing of successful pop-outs was performed by Plasmidsaurus using Oxford Nanopore Technology with custom analysis. For ΔsRNA_83 and ΔsRNA_24, no additional mutations were identified. For ΔsRNA_20, two additional point mutations were detected (C->G at position 50,650 in HVO_RS02135; A->G at position 1,284,523 in HVO_RS11505).

### Knockout phenotyping

To assess survival during acute oxidative stress, 3 replicates of WT and mutant strains were grown as described above in a total volume of 20 mL. 50 µL of culture were plated, and remaining culture was treated with H_2_O_2_. Cultures were plated on YPC with thymidine and incubated at 42°C for 4 days. Colonies were counted and used to calculate CFUs/mL and percent survival. To assess survival during chronic oxidative stress, 3 replicates of WT and mutant strains were grown as described above. Cultures were diluted into fresh media to an OD of 0.02 and treated with various H_2_O_2_ concentrations. Cultures were transferred to 100-well plates (250 µL/well) and grown in a Bioscreen C Pro for 42 hours at 42°C with shaking (normal speed, max amplitude, orbital). Absorbance measurements were taken every six hours. Data were plotted using R.

### qPCR

Samples for qPCR were prepared using 40 mL cultures from 3 replicates; half used for control samples and half treated with H_2_O_2_. Cells were harvested and RNA was extracted as described for RNA-seq library preparation. Conversion to cDNA was performed using a Superscript RT III kit (Invitrogen) with random hexamers. qPCR was conducted in 48-well plates with PowerUp™ SYBR™ Green Master Mix (Applied Biosystems, A25742) using the StepOne™ Real-Time PCR System with standard conditions and annealing temperature of 60°C (for primers, see Table S23). Fold change was determined by the Livak method (119).

## Data availability

RNA-seq reads, ribosome profiling reads, and annotation files are available on NCBI GEO: GSE293312 (RNA-seq) and GSE293326 (Ribo-seq). The mass spectrometry proteomics data have been deposited to the ProteomeXchange Consortium via the PRIDE (120) partner repository with the dataset identifier PXD062323 and 10.6019/PXD062323. Data analysis scripts are available on GitHub: https://github.com/diruggiero-lab/sRNAs.

## Acknowledgements

We thank Dr. Cesar Perez-Fernandez, Dr. Diego Gelsinger, and Frank Liu for experimental advice and helpful discussions. We thank Matthew F. Isada for extensive manuscript editing and helpful discussions. This work was supported by US National Science Foundation grants MCB-2034271 and MCB-57951. We would also like to thank the NSF Division of Molecular and Cellular Biology for a CAREER grant (Grant No. 2045844) and NIH/NIGMS for a New Innovator Award (No. DP2-GM140926) both to SDF. SDF further acknowledges support from the Camille & Henry Dreyfus Foundation and the Sloan Foundation. HMM is thankful for the Chemistry-Biology Interface Program training grant (Grant No. NIH-T32GM080189-13).

E.D.: Conceptualization, Methodology, Investigation – RNA-seq, Ribo-seq, mass spectrometry sample preparation, Western blots, knockout phenotyping, Formal analysis – analysis of sequencing data, footprint distribution analysis, Visualization, Writing – original draft, Writing – review & editing; H.M.M: Investigation – mass spectrometry sample preparation, mass spectrometry data acquisition, Formal analysis – mass spectrometry data analysis, Writing – review & editing; S.R.C: Formal analysis – DESeq2 analysis for all datasets; A.K.: Investigation – construction of tagged sRNA constructs and sRNA knockout strains, qPCR analysis; B.Y.H.H.: Investigation – construction of tagged sRNA constructs; S.D.F.: Methodology, Writing – review & editing; J.D.: Conceptualization, Methodology, Funding acquisition, Writing – review & editing, Supervision. All authors read and approved the final manuscript.

